# MHC Class I-like Gene in Squirrel Monkey Cytomegalovirus and the Evolution of the Virus

**DOI:** 10.1101/2025.04.10.648282

**Authors:** Eiichi Hondo, Tetsuya Mizutani, Hiroshi Shimoda, Atsuo Iida

## Abstract

Homology searches between viral and human peptides revealed that the S9 protein of squirrel monkey cytomegalovirus (*Saimiriine* herpesvirus 4 strain [SBHV4]) shares 60.6% amino acid identity across 86.3% of full-length human HLA-A. This represents the most extensive and highest degree of homology reported to date between a viral protein and an MHC class I molecule.

Phylogenetic analysis of MHC class I-like DNA sequences from primate HLA-A orthologs and SBHV4-S9 gene suggested that SBHV4-S9 was likely acquired via LINE-1-mediated horizontal gene transfer from the African Old World monkey *Piliocolobus tephrosceles* or a close ancestor 1.63-6.02 million years ago. SBHV4-S9 appeared to have undergone purifying selection following divergence from *P. tephrosceles*. Furthermore, phylogenetic analysis of the SBHV4-S1 gene and primate SLAMF6-like genes suggested that this gene was acquired after divergence from a close ancestor of the New World monkeys *Saimiri oerstedii* and *Saimiri sciureus*.

Comprehensive evaluation of the similarity between eight SBHV4 genes (including S1 and S9) and their respective homologs in 241 primate species, assessed via PCA-syncmer-based Wasserstein distances, revealed that the overall relatedness between SBHV4 and African *Cercopithecus ascanius*, *Cercopithecus pogonias*, and *Allenopithecus nigroviridis*, respectively was significantly greater (i.e., exhibited shorter Wasserstein distances) in each case than that observed with all examined members of the genus *Saimiri* and *Piliocolobus tephrosceles*. Collectively, these findings indicate that SBHV4, although isolated from *Saimiri sciureus* in South America, appears to have a historical connection with Old World monkeys in Africa, despite the typically high host specificity of cytomegaloviruses.

## Introduction

Vertebrate genomes harbor a diverse array of virus-like sequences [1]. These sequences represent remnants of ancient viral integrations resulting from host-virus interactions and exist along a continuum ranging from static accumulations to structurally and functionally dynamic elements. The distinction between static and dynamic states is likely influenced by the progressive accumulation of mutations over time rather than representing a clear dichotomy.

Endogenous retroviruses (ERVs), in particular, exhibit dynamic behavior. Including those rendered static through mutational inactivation, ERVs are estimated to constitute approximately 8% of the human genome [2]. Although implicated in host defense and cellular functions [3], ERVs have also been associated with the pathogenesis of diseases such as cancer and Alzheimer’s disease [4, 5]. Furthermore, retroelements such as SINEs have been implicated in the establishment of chromosomal domain boundaries in the mouse genome [6], suggesting that these elements have played critical roles in shaping and maintaining vertebrate morphology and function from ancient times to the present.

The substantial proportion of retroelements in vertebrate genomes, evidence of recent retroviral endogenization [7], and the gradation of sequences from static to dynamic collectively render retroelements an intriguing subject of investigation. While retroviruses possess enzymes that facilitate the integration of their genomes into host DNA, the integration of other viral genomes is currently understood to depend largely on stochastic events. Consequently, intact and transcriptionally active non-retroviral sequences are rarely found in vertebrate genomes. It is therefore inferred that ancient ancestral viruses integrated into host genomes and now persist in a static state or as quasi-dynamic elements with limited transcription in extant vertebrates. Nevertheless, even static sequences, when present in large copy numbers, have the potential to influence higher-order chromatin architecture, and transcriptional activity originating from such non-retroviral elements may modulate immune responses during infection with related viruses.

Although early compilations of non-retroviral integrations date back to the early 2000s [1], recent increases in virome-scale datasets and advances in bioinformatics have enabled more comprehensive analyses [8]. A previous study identified 2,040 virus-like sequences within 295 vertebrate genomes by screening 24,478 viral proteins from the Shotokuvirae (16 families) and Orthornavirae (112 families) kingdoms (tblastn E-value 1e-5), and discussed host phylogenetic relationships across major viral groups.

In the present study, a search was performed using the FASTA program [9] against the complete set of peptides derived from the human genome build GRCh38, employing all viral proteins registered in the Virus-Host DB (RefSeq; 1,860,785 entries as of 2024; https://www.genome.jp/virushostdb/). This analysis revealed a high degree of homology between the S9 protein of Simian beta-herpesvirus 4 (simian cytomegalovirus; SBHV4) and the human Major Histocompatibility Complex class I molecule HLA-A, with coverage spanning 86.3% of the full-length HLA-A protein. Notably, this S9 protein is unique and was identified in a genomic region distinct from previously reported herpesviral regions detected in comprehensive searches for virus-like proteins [8]. In this study, the evolution of SBHV4 is discussed, with a focus on several proteins including S9, from a perspective distinct from the hypothesis of ancient endogenization of a *Saimiriine* beta-herpesvirus ancestor in the human lineage.

## Results … Discussion

Homology searches using FASTA36 software between the entire viral peptide set and the human peptide set yielded 372 hits with ≥60% identity and ≥200 amino acid residues. Among these, we focused on a 315-residue overlap (86.3% of full-length HLA-A) between the membrane protein S9 (SBHV4-S9; 344 residues in length) of *Saimiriine* herpesvirus 4 strain (NC_016448.1) and human major histocompatibility complex class IA (HLA-A; 365 residues in length), showing 191 identical amino acids (60.6%) (Supplemental data 1). This represented the most extensive and highest homology observed thus far among viral proteins and an MHC class I molecule.

Further homology analysis of the SBHV4-S9 peptide, restricted to the HLA-A homologous region, against all other known viral peptides revealed no substantial similarity, with the highest identity being 43.2% (137/317) to a sequence from *Murmansk poxvirus* (Supplemental data 2). SBHV4-S9 exhibited higher similarity to mammalian peptides than to any viral sequence, underscoring its unique status among viral proteins. Given the low probability that the corresponding proteins in the viral species listed in Supplemental data 2 shared a recent common viral ancestor, phylogenetic analysis was not performed.

Structural analysis was conducted on the full-length peptide sequences of SBHV4-S9 and HLA-A. Based on subsequent phylogenetic analysis, we included the MHC class I-like gene from *Piliocolobus tephrosceles*, the closest relative to SBHV4 (Fig. 4), and that from *Scleropages formosus*, the most distantly related (Fig. 3), for structural comparison (Fig. 1). Visual inspection revealed a high degree of structural similarity among the four (Fig. 1, upper panel; SBHV4-S9 magenta, HLA-A cyan, *P. tephrosceles* gold, *S. Formosus* green). Template Modeling (TM)-scores and Root Mean Square Deviation (RMSD) values between SBHV4-S9 and HLA-A, SBHV4-S9 and “MHC class I-like” (*P. tephrosceles*), and SBHV4-S9 and “MHC class I-like”(*S. formosus*) were (0.83, 1.98 Å), (0.89, 1.98 Å), and (0.78, 2.86 Å), respectively, further demonstrating significant structural resemblance based on intermolecular distances. The SBHV4-S9 peptide exhibited greater similarity to HLA-A and the *P. tephrosceles* MHC class I-like peptide than to the *S. formosus* MHC class I-like peptide. Superimposition of the SBHV4-S9 and HLA-A structures showed particularly high similarity in the α1, α2, and α3 domains of HLA-A, with SBHV4-S9 also possessing the peptide-binding pocket between α1 and α2. However, differences in the angles at the base of the branching towards the N- and C-termini and variations in individual molecular features were observed (Fig. 1 A and B). Considering the potential for cross-species infection, the immunogenicity of this structure itself warrants consideration. The human cytomegalovirus UL18 protein has been previously reported as the viral protein exhibiting the highest peptide homology to HLA, displaying approximately 25% sequence identity [10]. UL18 has been shown to bind inhibitory NK cell receptors with surprisingly high affinity (over 1000-fold greater than HLA), suggesting a potential mechanism for NK cell function suppression [11].

**Figure 1.**
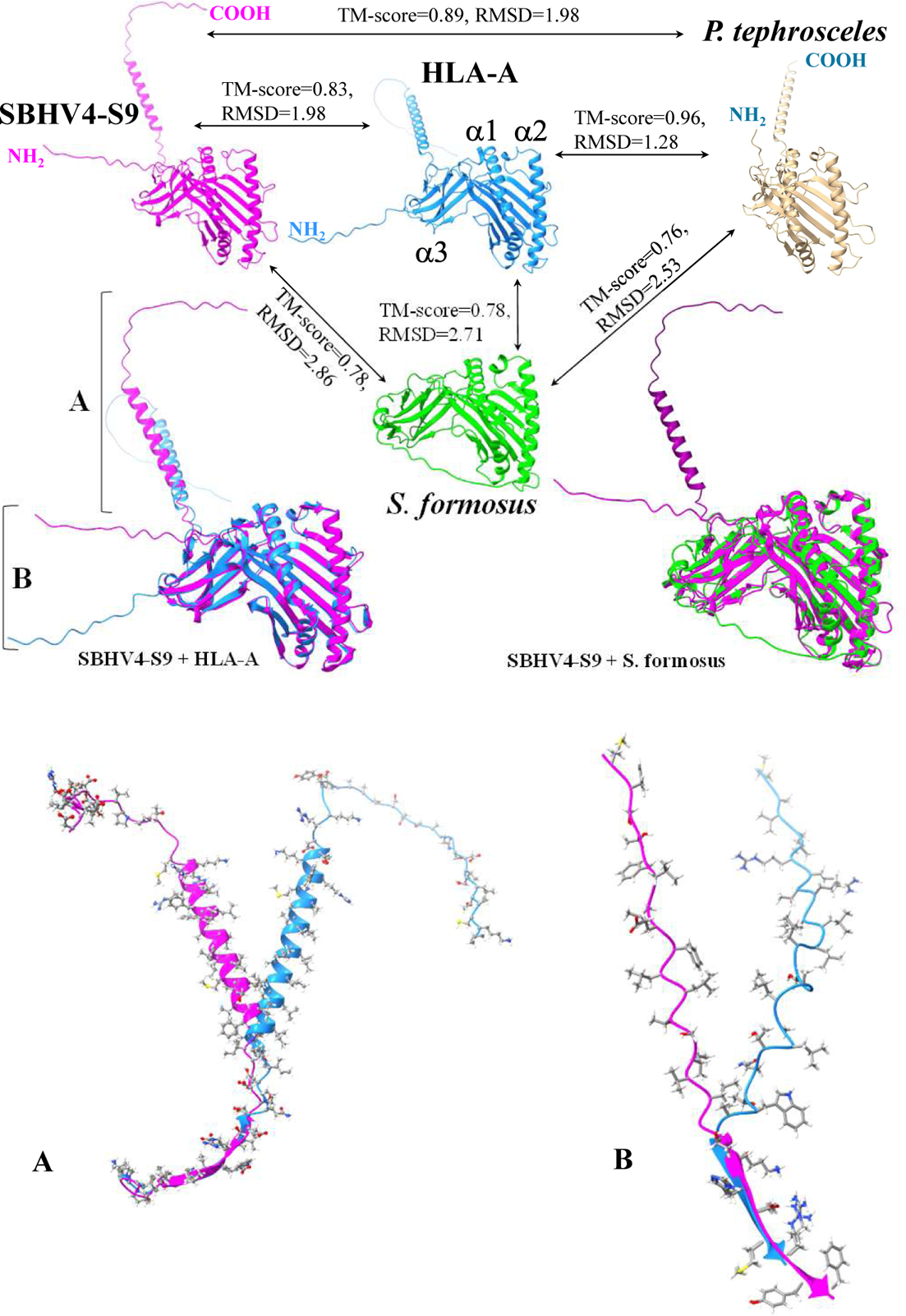
Molecular Models of MHC Class I-like Peptides (AlphaFold). Magenta: SBHV4-S9; cyan: human HLA-A; gold: *Piliocolobus tephrosceles* MHC class I-like peptide; green: *Scleropages formosus* MHC class I-like peptide. α1, α2, α3 denote the HLA-A subunits. **Middle Left:** Superimposed structures of SBHV4-S9 and HLA-A. **Middle Right:** Superimposed structures of SBHV4-S9 and *Scleropages formosus* MHC class I-like peptide. TM-score and RMSD values quantifying structural similarity between the respective molecular models are indicated. **Bottom:** Structural divergence between SBHV4-S9 (magenta) and HLA-A (cyan) at the C-terminus (A) and N-terminus (B).

What are the implications of the uniqueness of the SBHV4-S9 protein among previously isolated or sequenced viruses, and its striking structural similarity to HLA-A? Beyond the integration of viral genomes into the host genome, as observed with endogenous retroviruses, there are several documented instances suggesting the acquisition of host genomic fragments by viruses. For example, in the *Pestivirus* genus, the genomes of three bovine viral diarrhea virus (BVDV) strains exhibit distinct insertion sites of host-derived ubiquitin genes. Notably, the pathogenic CP1 strain and the non-pathogenic NCP1 strain, isolated from the same bovine host, showed the presence of a host ubiquitin gene insertion specifically in CP1, which was absent in NCP1. The potential role of this ubiquitin gene insertion in the differential pathogenicity has been discussed [12].

Furthermore, studies on poxviruses, which are DNA viruses, have revealed the presence of target site duplications (TSDs) generated by the host Long Interspersed Nuclear Element-1 (LINE-1) retrotransposon. This observation suggests a LINE-1-mediated molecular mechanism for the incorporation of host genomic fragments (DNA reverse-transcribed from mRNA) into the viral genome [13].

Regarding the squirrel monkey cytomegalovirus focused on in this study, its S1 protein (285 amino acids; NC_016448.1) has been shown to possess high homology to the signaling lymphocytic activation molecule (SLAM) family receptors of the Bolivian squirrel monkey (77% identity to sbSLAMF6) [14]. It has been proposed that this S1 protein was acquired through retrotransposition during the co-evolution of the virus and its host. Similarly, the absence of intron sequences characteristic of HLA-A within the homologous region of SBHV4-S9 may indicate a potential acquisition via reverse transcription. This specific S1 homologous sequence was also identified in our current investigation (Supplemantal data 1).

Initially, we conducted a search for TSDs flanking the SBHV4-S9 gene. Multiple sequences, suggestive of TSDs, were found at both ends of the HLA-A homologous region within the S9 gene, as well as at both ends of the entire S9 gene (Fig. 2 The blue gradient boxes in the upper panel). While TSDs mediated by LINE-1 are typically directly adjacent to the inserted DNA, the presence of pseudo-empty sites containing polyadenylation signals can serve as supplementary evidence for LINE-1 involvement [13]. In this study, some clear polyadenylation signals were observed within the low GC content regions; however, due to accumulated mutations, the authentic polyadenylation signal utilized remains undetermined. Consequently, while the precise length and integration site of the SBHV4 MHC class I-like sequence cannot be definitively established, these findings suggest a potential transfer of genetic material from the host genome to this cytomegalovirus via a LINE-1-mediated mechanism, analogous to that reported in poxviruses.

**Figure 2.**
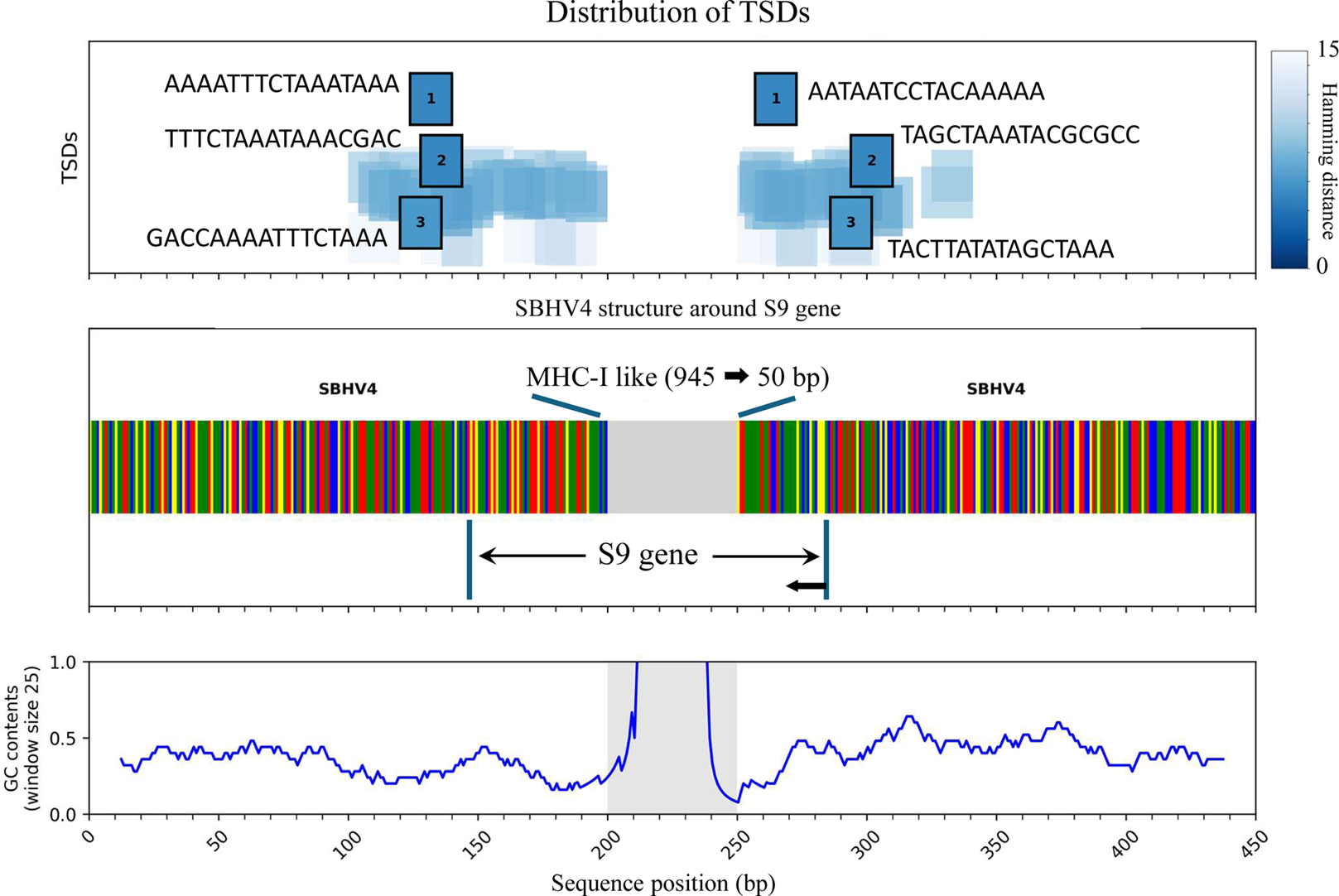
Predicted Target Site Duplications (TSDs) Flanking the SBHV4 S9 Gene. The middle color-coded section illustrates the MHC class I like gene in S9 gene (grey) and surrounding nucleotides (Each nucleotide: adenine (A) in green, thymine (T) in red, guanine (G) in yellow, and cytosine (C) in blue). For visualization, the 945 bp inside S9 gene is compressed to 50 bp (referred to as MHC-I like). Putative flanking short sequences, suggestive of TSDs, are observed at both ends of the compressed HLA-A homologous region in and around the S9 gene (the upper panel). The top three pairs with the shortest Hamming distances were numbered within the box, with the complete sequence for each pair listed. The bottom panel displays a histogram (window size 25) of the GC content (A value of 1.0 corresponds to 100% GC contents.).

Phylogenetic analysis (maximum likelihood (ML) method) was performed on a total of 133 MHC class I-like gene sequences, including the finally obtained SBHV4-S9 and seven human variants. To initially ascertain the branching position of SBHV4-S9, a cladogram devoid of genetic distance information was generated to assess its placement and statistical support (Fig. 3). Both SHaLRT and Ultra-fast bootstrap (UFbootstrap) values by IQ-TREE 2 were calculated, with values ≥80% and ≥95%, respectively, considered reliable. As no integrated index for assessing both measures was available, a new index, the S-value, based on their geometric mean, was calculated, with a threshold of 87.0 established for reliable support 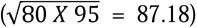. Phylogenetic analysis (ML) using *S. formosus* as an outgroup confirmed robust branching leading to SBHV4-S9, as depicted in Fig. 3. To conduct a more detailed phylogenetic analysis (ML method) of the SBHV4-S9 gene, which had diverged from a primate clade (closest to *Papio anubis*), we incorporated more primate species into our dataset. Analysis of 186 primate species together with the SBHV4-S9 gene revealed that SBHV4 diverged from *P. tephrosceles* after the split among Old World monkeys (Fig. 4). The topology of the clade comprising 11 Old World monkey species containing the SBHV4-S9 gene was consistent with that inferred for the same clade, excluding SBHV4, by Bayesian inference (Fig. 5, right panel). For this clade, both the ML S-values and the Bayesian posterior probabilities (BPP) were high for all nodes. The genetic distance between SBHV4-S9 and *P. tephrosceles* estimated by ML was substantial (Fig. 4, right panel). Bayesian inference was performed excluding SBHV4-S9 for the following reasons: initial analysis including SBHV4-S9 placed its divergence basal to Asian arowana (*S. formosus*), a discrepancy likely arising from the substantial post-divergence distance of SBHV4-S9 as indicated by ML analysis. Viruses can undergo rapid diversification during specific short periods, potentially driven by interspecies transmission and spread [15], a phenomenon likely experienced by SBHV4, distinct from vertebrate evolution. The high sequence similarity of SBHV4-S9 to primate peptides revealed by ML analysis (Fig 4, right panel) and homology searches (Asian arowana (*S. Formosus*) was ranked outside the top hits in Supplemental data 3.), coupled with the greater structural similarity of SBHV4-S9 peptides to HLA-A and *P. tephrosceles* MHC class I-like peptides compared to that of Asian arowana (Fig. 1), supported the exclusion of a pre-arowana divergence for SBHV4-S9. Consequently, Bayesian inference was conducted without the SBHV4-S9 sequence. The divergence time among the other 10 Old World monkey species belonging to the same clade as *P. tephrosceles* was considered the oldest likely divergence of SBHV4-S9 from *P. tephrosceles* (Fig. 5, right panel; red arrow), estimated to have occurred between 1.63 and 6.02 million years ago (Height: 3.68 million years) (Fig. 5, right panel; violet arrow). This suggests that SBHV4-S9 likely diverged from *P. tephrosceles* or a closely related ancestor within this timeframe. However, the extended branch length observed for SBHV4-S9 in the ML phylogeny suggests a far older time to the most recent common ancestor than this range. This discrepancy should be attributable to the viral mutation rate being substantially accelerated and entirely independent of that of the host MHC-class I genes.

**Figure 3.**
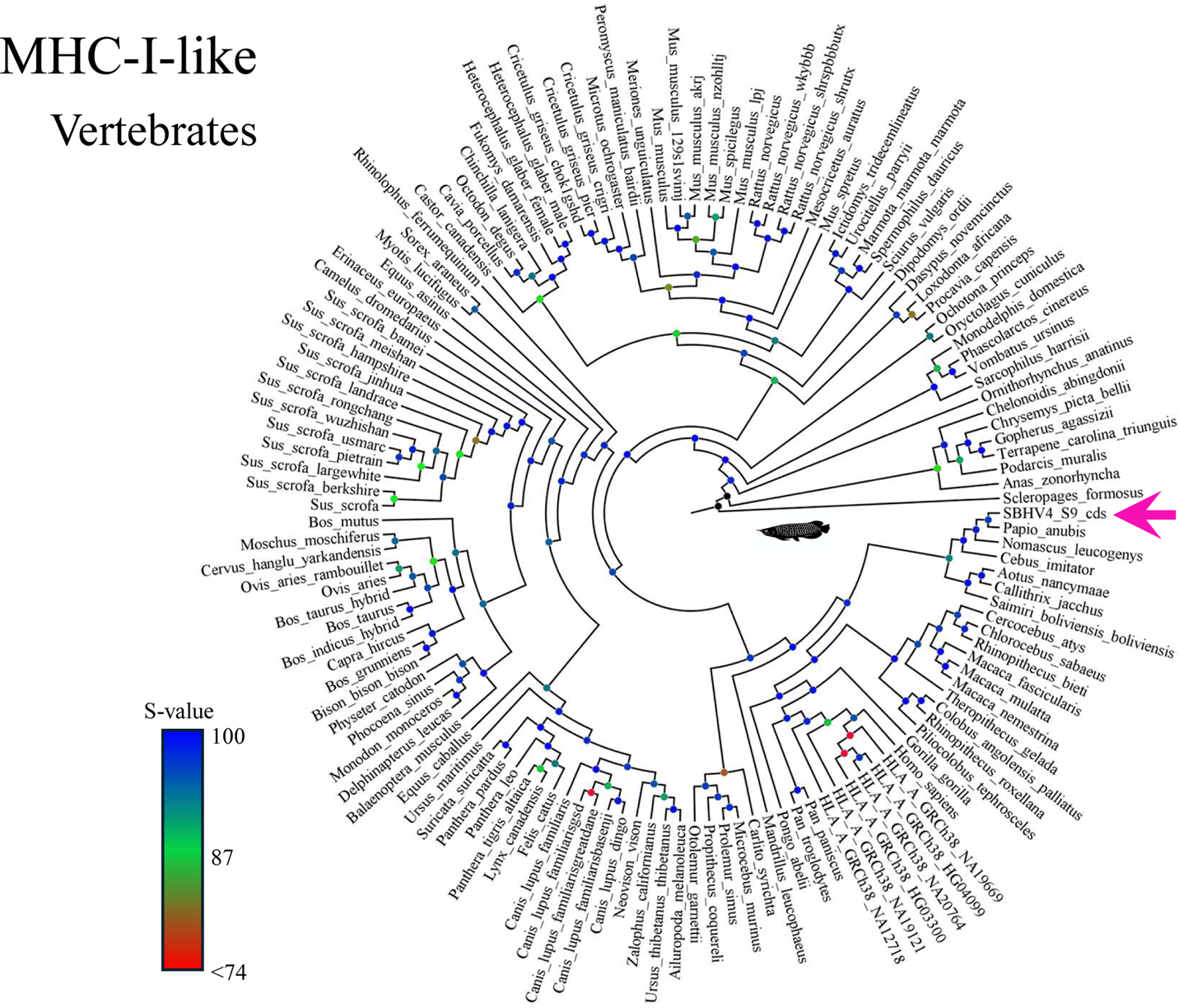
Phylogenetic analysis of the SBHV4-S9 gene among vertebrate MHC class I-like sequences. Maximum-likelihood cladogram depicting the branching patterns of 133 FASTA sequences, including SBHV4-S9 and seven human individuals. The position of SBHV4-S9 is indicated by a magenta arrow. Node colors denote S-values (the geometric mean of SH-aLRT and UFbootstrap values), with values ≥87 considered to provide reliable support.

**Figure 4.**
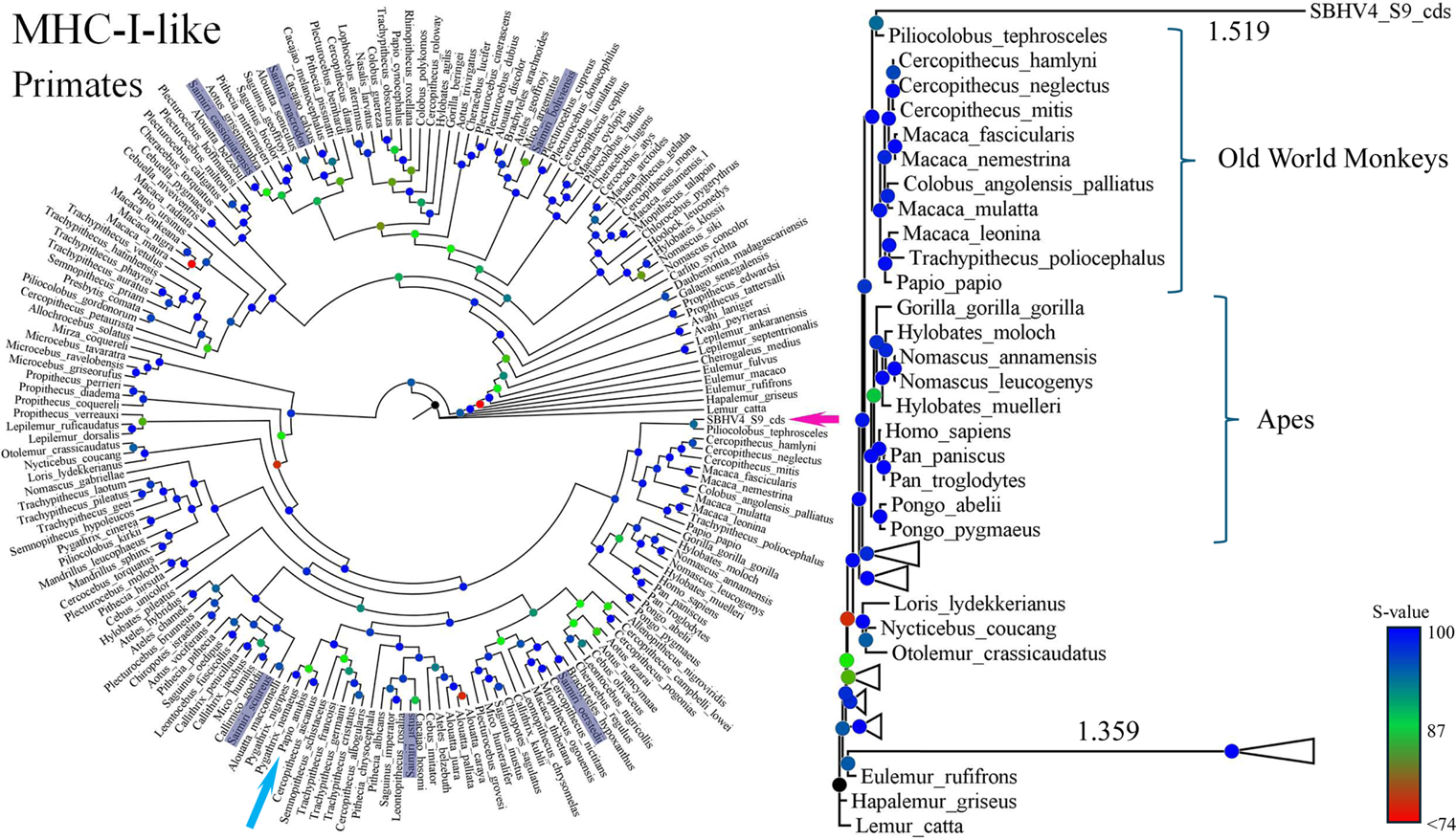
Phylogenetic relationships between MHC class I-like genes from 186 primate species and the SBHV4-S9 gene. Left panel: Cladogram inferred by the maximum-likelihood method, omitting genetic distances. The position of SBHV4-S9 is indicated by a magenta arrow. Species belonging to the genus Saimiri are highlighted in the species names. A cyan arrow denotes *Papio anubis*, whose phylogenetic placement represents the closest relationship to SBHV4-S9 among vertebrate species, as shown in Fig. 3. Node colors reflect the magnitude of S-values. *Right panel:* A maximum likelihood phylogenetic tree incorporating genetic distances. The SBHV4-S9 gene diverges from *Piliocolobus tephrosceles* with an extended branch length.

**Figure 5.**
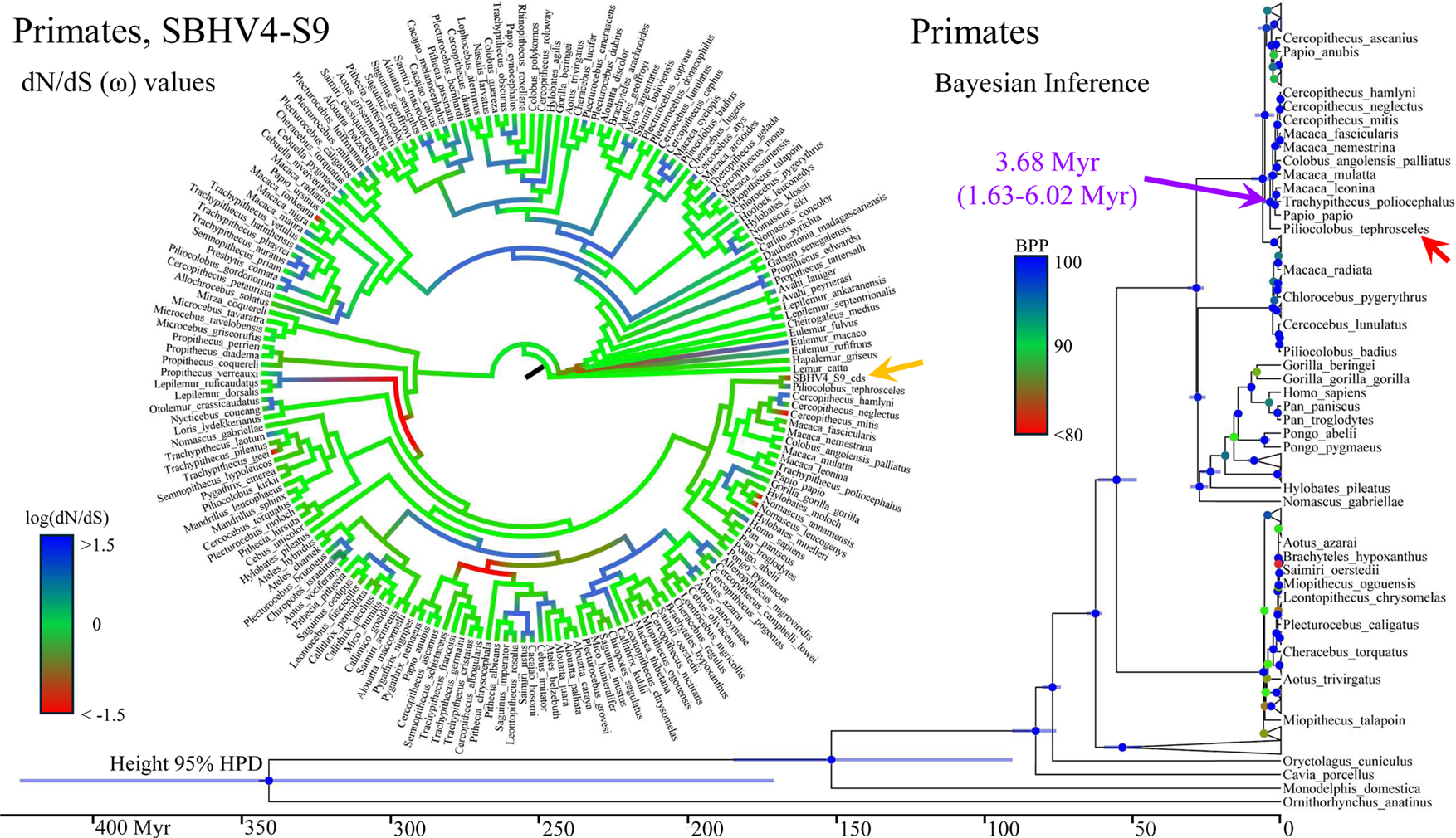
dN/dS (ω) Values and Bayesian Phylogenetic Tree. **Left:** dN/dS (ω) values for each branch calculated from 186 primate sequences. The yellow arrow indicates the SBHV4-S9 gene. Values ≥1.5 (log□□(ω)) are red, 0 (log□□(ω) = 1) is green, and ≤ −1.5 (log□□(ω)) are blue. Purifying selection is inferred for SBHV4-S9. **Right to Bottom:** Bayesian phylogenetic tree excluding SBHV4-S9. The purple arrow indicates the divergence point of *Piliocolobus tephrosceles*. Node colors represent Bayesian Posterior Probabilities (BPP; threshold ≥80%): red (<80%), green (90%), blue (100%). Bars on nodes indicate 95% Highest Posterior Density intervals (HPD).

Collectively, one hypothesis posits that SBHV4-S9 was transferred from the *P. tephrosceles* MHC class I-like gene to SBHV4 via LINE-1, approximately 1.63 to 6.02 million years ago. The current host range of SBHV4 includes the common squirrel monkey *(Saimiri sciureus)*, a New World primate, while *P. tephrosceles* is an Old World primate, implying cross-species transmission across continents. To investigate the evolutionary dynamics of this process, we first calculated dN/dS (ω) values (Fig. 5, left panel), which were generally neutral across vertebrates. Although strong purifying selection or mild positive selection was observed in some primates, the values were generally consistent with neutrality. In SBHV4-S9, purifying selection was inferred. To assess this, a likelihood ratio test (LRT) was conducted using SBHV4-S9, yielding an LRT statistic of 283.11 (df = 1), *p* < 1.59 × 10□^63^. In 10 million simulation trials, no instance exceeded the observed LRT statistic ( *p* < 1.0 × 10□□). These results rejected the null hypothesis and supported the hypothesis that SBHV4-S9 is under purifying selection. While purifying selection is common in viruses [16], our calculation was performed within the context of primate MHC class I-like sequences, not solely among viral strains. Given that the structure of the S9 protein is highly similar to that of the MHC class I molecule, especially its antigen-binding site (Fig. 1), it is plausible that purifying selection has acted on the S9 gene to maintain its function.

*P. tephrosceles* is a primate species native to central Africa, while SBHV4 was isolated from squirrel monkeys inhabiting the rainforests in Northern South America. Direct transmission of a virus with the highest sequence similarity to MHC class I like gene (*P. tephrosceles*) across such geographical barrier is improbable. Parallels could be drawn to the human introduction of yellow fever virus from Africa to South America [17]; however, historical records of human-mediated transport of *P. tephrosceles* to South America are lacking. It has been suggested that the *Chlorocebus sabaeus* was introduced from Africa to the West Indies in the 16th century, where it remains extant, while the habitat of *P. tephrosceles* was different [18].

Our hypothesis posits the transfer of an orthologous sequence to MHC class I like gene from *P. tephrosceles* to SBHV4. As previously noted, such instances of host gene acquisition by viruses are not unprecedented; as seen with the S1 protein of squirrel monkey cytomegalovirus is proposed to have originated from a SLAMF receptor gene of the Bolivian squirrel monkey [14]. Thus, supporting both our current hypothesis and previous findings suggests the potential for squirrel monkey cytomegalovirus to have acquired coding sequences from other primate genomes as well.

We performed ML phylogenetic inference between the SBHV4 S1 gene, previously suggested to have originated from the squirrel monkey SLAMF6-like gene, and primate (121 species) SLAMF6-like genes (Fig. 6). The analysis confirmed that the S1 gene diverged from *Saimiri boliviensis* (and possibly from the common ancestor of *Saimiri oerstedii* and *Saimiri sciureus*).

**Figure 6.**
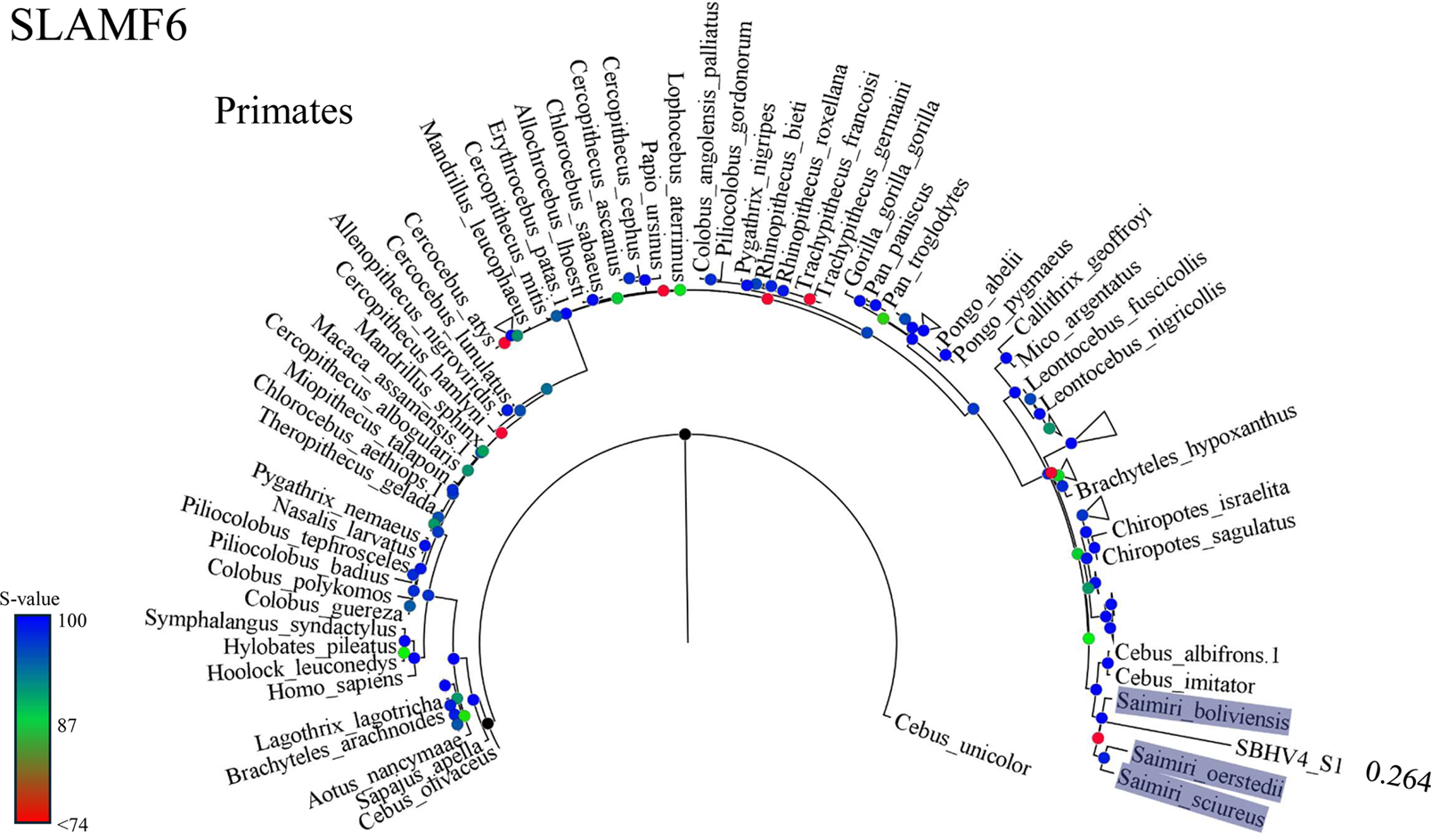
Maximum Likelihood Phylogenetic Analysis of 121 Primate SLAMF6-like and SBHV4-S1 Genes. Node colors represent S-values (geometric mean of SHaLRT and UFbootstrap values). SBHV4-S1 arose from a lineage that had already separated from *Saimiri boliviensis*, and it later branched from the ancestor of *Saimiri oerstedii* and *Saimiri sciureus*.

**Figure 7.**
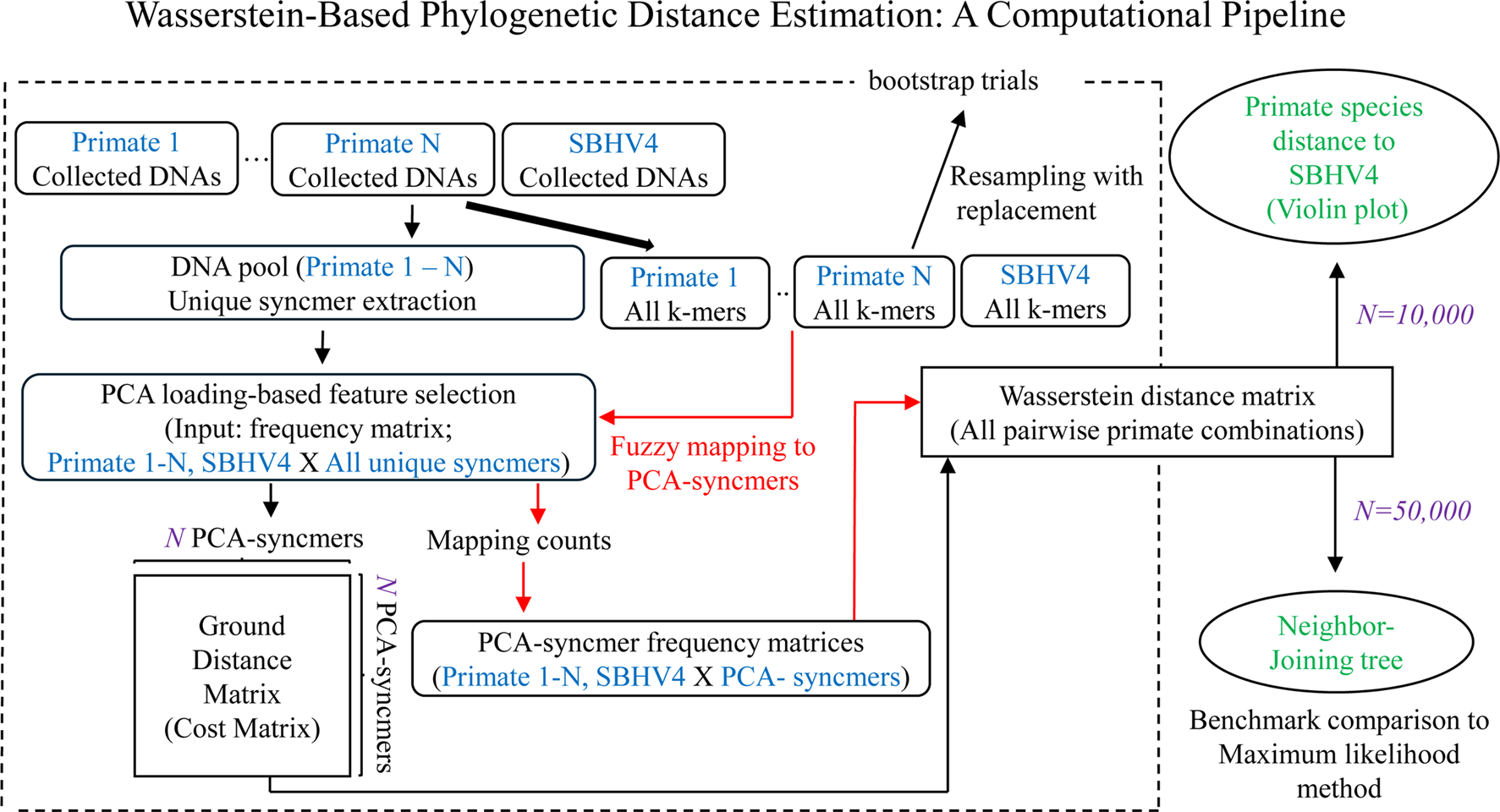
Wasserstein-Based Phylogenetic Distance Estimation: A Computational Pipeline. The diagram illustrates the integrated alignment-free framework for inferring phylogenetic relationships using the Wasserstein distance. The symbol *N* represents the number of representative PCA-syncmers used: *N* = 10,000 for the primary SBHV4 analysis and *N* = 50,000 for the benchmarking analysis (as detailed in the Materials &Methods). The overall process involves three main stages: (1) Feature Selection: Initial k-mer frequency vectors are generated after unique syncmer extraction, from which *N* representative PCA-syncmers are chosen via PCA loading-based feature selection for dimensionality reduction. (2) Cost Matrix Construction: An N × N ground distance matrix is computed based on weighted Hamming distances between these selected PCA-syncmers, serving as the cost matrix input for Wasserstein distance calculation. (3) Final Frequency Matrices (Vectors) (Mass Distribution) Construction: K-mer sets from each species (query mass) are fuzzy-mapped onto the PCA-syncmer feature space (reference feature space) using weighted Hamming distances and a similarity threshold. When a k-mer maps successfully, the corresponding syncmer count (“Mapping counts”) is incremented by a weight equal to the observed nucleotide similarity (e.g., 0.98 for 98% similarity). This generates L1-normalized PCA-syncmer frequency matrices for all species, which were used as Final Frequency Matrices. The Wasserstein distance matrix is then computed from these inputs, allowing for the reconstruction of phylogenetic trees. The pipeline also incorporates bootstrap trials to assess stability and includes a benchmark comparison to Maximum Likelihood methods using mitochondrial genes.

Based on the analyses conducted thus far, it appears reasonable to infer that SBHV4 has successively acquired several genes at different evolutionary stages, likely accompanying host transitions. On the other hand, it remains of considerable interest to determine which primate lineage, overall, exhibits the closest genetic affinity to the present SBHV4 genes. To address this, we employed a newly developed alignment-free analytical framework to comprehensively investigate the homologs of all SBHV4 genes across 241 primate species, with the aim of identifying which primates show the greatest overall similarity to those of SBHV4.

Specifically, for each SBHV4 protein, we retrieved all homologous genes from 241 animal species using Spaln3, and then calculated the Wasserstein distances between SBHV4 genes and their homologs in each primate species. Prior to the main analysis, the validity of this method was assessed through a benchmark comparison with ML test, after converting the Wasserstein distance matrices to newick format files by neighbor-joining (NJ) method. We used 49 species spanning tarsiers, New World monkeys, Old World monkeys, and apes, whose species phylogenetic relationships are well established [35, 36], and analyzed 13 mitochondrial DNA genes plus two rRNA sequences. For IQ-TREE 2, sequences were aligned for each gene across the 49 species using MAFFT, concatenated, and analyzed under a partitioned model to account for differences in substitution rates. Because Wasserstein distance matrices calculated here cannot be directly compared with ML outputs, we applied NJ clustering to the Wasserstein distance matrices, generated a newick format file, and visualized the resulting trees for comparison (ML estimation is not possible directly from distance matrices) (Fig. 8). Correlation between pairwise path lengths from the NJ trees and the Wasserstein distances was assessed using a Mantel test, revealing a highly significant correlation (r = 0.9657, P = 0.0001). This indicates that pairwise path lengths from NJ clustering can serve as a proxy for Wasserstein distances. While the ML analysis did not preserve known phylogenetic relationships among the tested species (Fig.8 left panel), the present method retained the major clade separations (Fig. 8 right panel): divergence of tarsiers from the primate ancestor, that of Genus *Aotus*, followed by the split between Genus *Saimiri* and the lineage leading to Old World monkeys and Apes. Within the apes, the method correctly placed gibbons as diverging from the common ancestor of the Hominoidea, but subsequently grouped humans with gorillas, deviating from the established species-level topology [35, 36]. Nonetheless, in the broader context, this multi-gene phylogenetic analysis exhibited performance higher to that of ML test in this mitochondrial samples, supporting the robustness of our approach using multiple genes.

**Figure 8.**
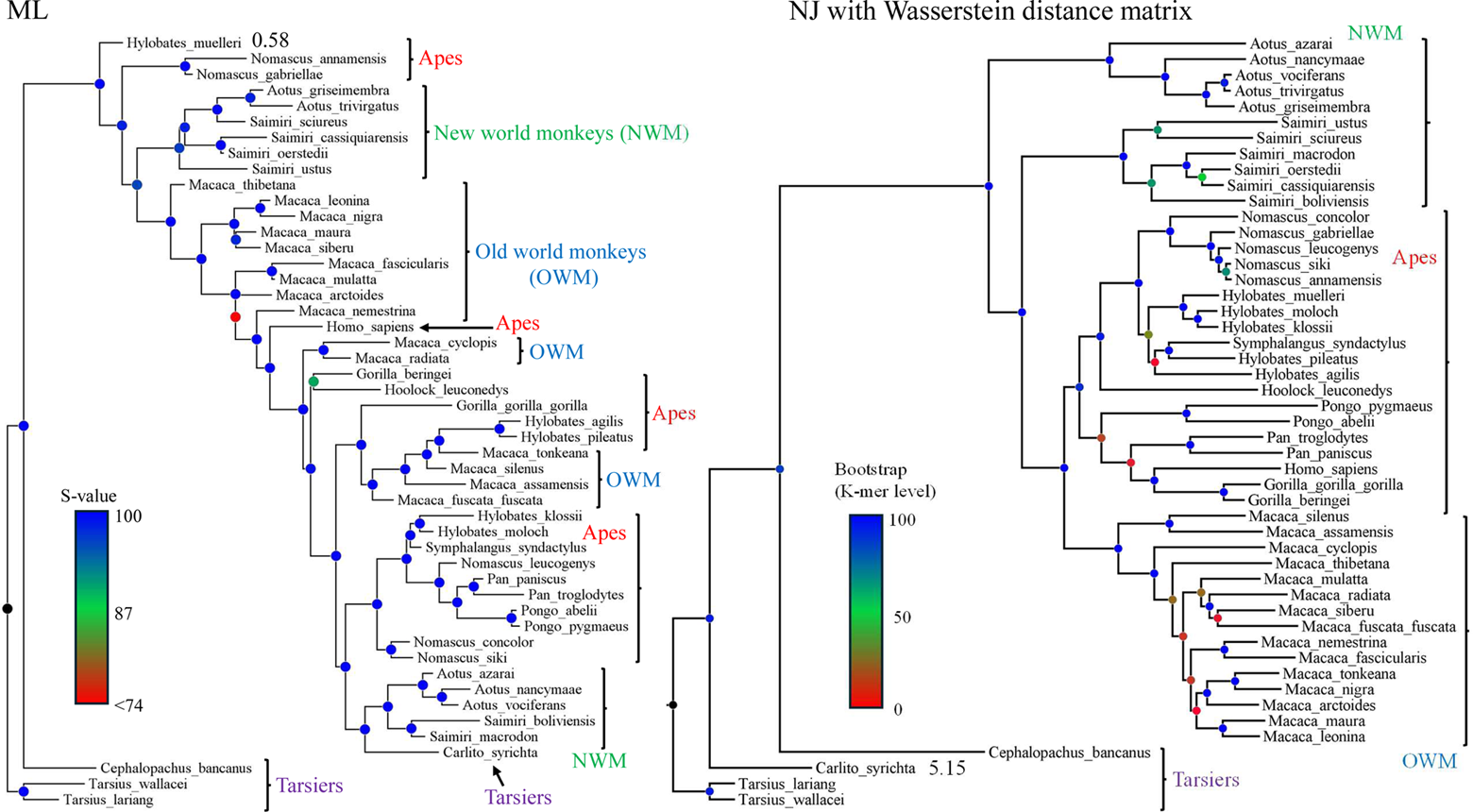
Benchmark comparison between phylogenetic trees inferred using the Neighbor-Joining method based on Wasserstein distances and the maximum likelihood method. The ML tree (left panel) was inferred using IQ-TREE 2 with 15 mitochondrial and rRNA genes (13 protein-coding, 2 rRNA) from 49 primate species, including Tarsiers, New World Monkeys (NWM), Old World Monkeys (OWM), and Apes. Genes were individually aligned with MAFFT, concatenated by species, and partitioned for the IQ-TREE 2 analysis. Node support values are represented by Sh-aLRT (S-values). The NJ tree (right panel) was constructed from the Wasserstein distance matrices. Bootstrap support values were calculated using 200 kmer-level resamplings with replacement (200 Wasserstein distance matrices). While this NJ tree does not fully recapitulate the currently accepted species phylogeny, the overall topology, separating Tarsiers, followed by NWM, and then OWM and Apes, is retained, despite some discrepancies observed at the genus and species levels compared to established phylogenetic trees.

We used violin plots (300 k-mer-level bootstrap trials) to visualize the Wasserstein distances between SBHV4 and 241 primate species, respectively. These plots highlighted two key points: the ten species exhibiting the shortest distances (Fig. 9, cyan), and those exhibiting significantly longer distances to the ten species (Fig. 9, brown, based on Bonferroni-corrected Dunn’s post-hoc test). Interestingly, the significantly longer-distance species included all members of the genus *Saimiri* examined (despite some showing the highest relatedness in the SBHV4 S1 and SLAMF6-like genes) and *P. tephrosceles* (which harbors the most closely related MHC-I-like gene to SBHV4 S9 gene). While six of the top ten species consisted of New World monkeys, Old World monkeys (Cercopithecus ascanius, Allenopithecus nigroviridis, and Cercopithecus pogonias) and one lemur (Propithecus deckenii coronatus) were also part of this group.

**Figure 9.**
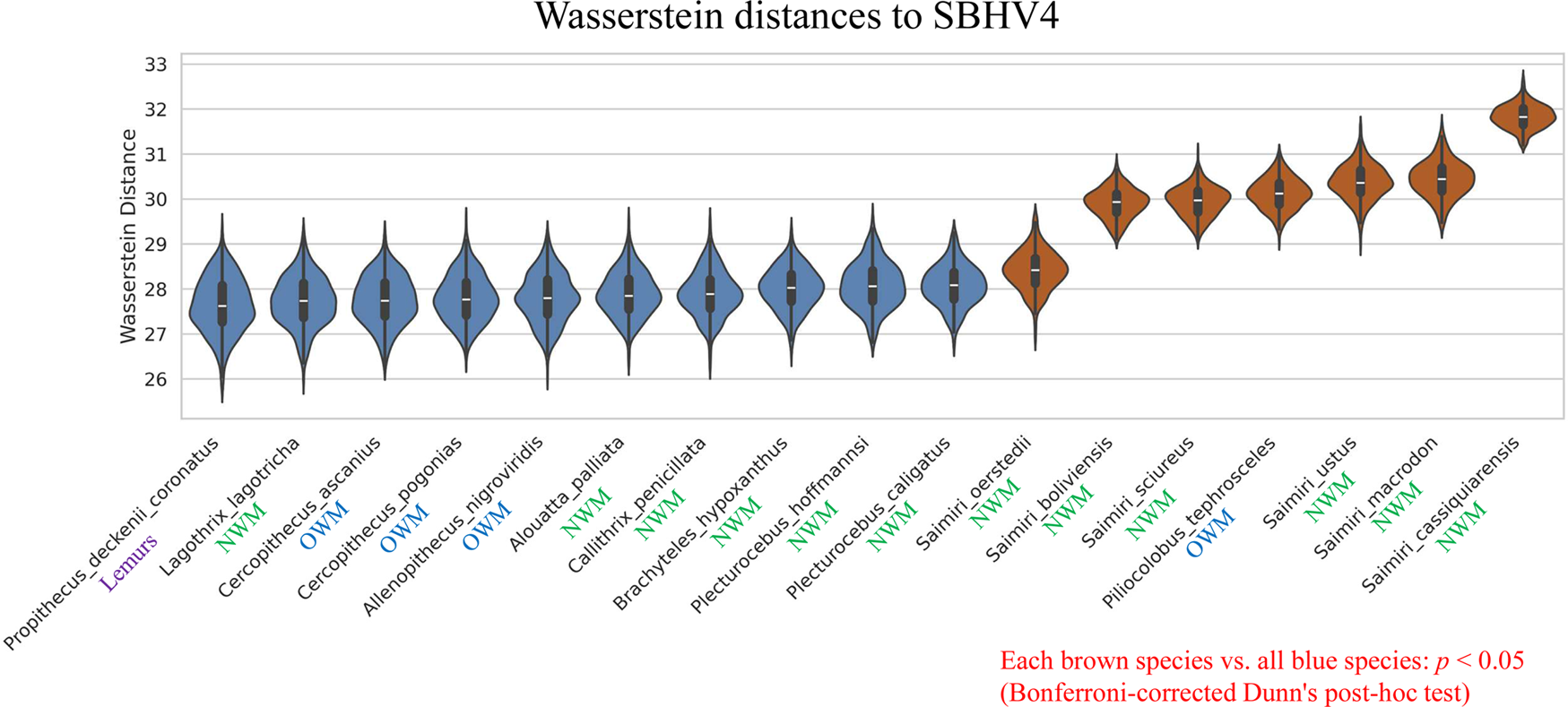
Comparison of Wasserstein distances to SBHV4. Violin plots show the Wasserstein distances between SBHV4 and 17 primate species. The cyan plots on the left represent the ten species exhibiting the shortest distances (closest to SBHV4), while the brown plots on the right represent the seven species exhibiting significantly longer distances (more distant to SBHV4) (Significant difference was found between each brown violin plot and all blue violin plots by Bonferroni-corrected Dunn’s post-hoc test, p < 0.05). The taxonomic group (Lemurs, OWM: Old World Monkeys, NWM: New World Monkeys) is indicated below each species name.

All primates listed in Fig. 9 had only S1 and S9 homologs. Other homologs may have contributed to an increased Wasserstein distance to SBHV4, which suggests a lower degree of relatedness to SBHV4 for these other homologs compared to the S1 and S9 homologs. Importantly, our current methodology showed that overall SBHV4 genes were more closely related to some Old World monkeys than to the genus *Saimiri*, as was observed between the SBHV4 S9 gene and the MHC class I-like gene. Furthermore, the reason for the emergence of *P. deckenii coronatus*, a species endemic to Madagascar, as being closer to SBHV4 than the genus Saimiri in the current analysis is yet to be elucidated.

Although SBHV4 was originally isolated from *S. sciureus*, and despite the generally strict host specificity of cytomegaloviruses, the data suggest a historical capacity to infect genetically distant hosts obtaining host genes. At present, the origin of SBHV4 cannot be conclusively assigned to either South America or Africa. Nevertheless, the results of this study indicate a deeply rooted association between SBHV4 and African primates.

## Materials&Methods

### Entire Viral Peptide vs. Entire Human Peptide Set

The complete viral peptide set (release 225), updated August 6, 2024, was downloaded from GenomeNet (Virus-Host Database). The entire human peptide set, corresponding to the GRCh38 genome build, was obtained from NCBI. Homology searches were performed using FASTA36 (-E 1e-20) [9], and results were screened for hits with ≥60% identity and ≥200 amino acid residues (Supplemental data 1).

### SBHV4-S9 vs. All Viral Peptides

To investigate the relationship of SBHV4-S9 (YP_004940188.1; 344 residues, full length) to other viral peptides, a FASTA36 homology search was performed using the SBHV4-S9 peptide sequence (HLA-A homologous region only) and the entire viral peptide dataset (Supplemental data 2).

### SBHV4-S9 TSD Identification

To investigate the presence of TSDs flanking the HLA-A homologous region of SBHV4-S9, TSD searches were performed using the SBHV4 genome (NC_016448.1) and the region exhibiting high HLA-A homology to SBHV4-S9. The SBHV4-S9 locus within the SBHV4 genome was identified, and 200 bp upstream and downstream flanking sequences were retrieved. The 5′ flanking region was split into overlapping fragments using a 4-20 bp sliding window, and each fragment was used as a query against the 3′ flanking region using the Needleman-Wunsch alignment algorithm [19] (Code 1; The output files TSD_plus.txt and TSD_minus.txt present the results of two complementary approaches to identifying the Target Site Duplication (TSD) flanking the insertion element (IE) in the reference genome. They essentially represent the search from opposite directions of the insertion site.). TSD candidates exhibiting ≥60% identity were exported in FASTA format, converted to FASTQ format (assigned with maximum quality scores), and subsequently mapped using HISAT2 [20].

### SBHV4-S9 vs. Peptides from Other Animal Species

To identify peptide sequences homologous to SBHV4-S9 in diverse species, the SBHV4-S9 peptide (HLA-A homologous region only) was queried against the comprehensive peptide datasets of all species available at https://ftp.ensembl.org/pub/release-112 using a homology search (-b 1 -d 1 -E 1e-10 -m BB). Results were filtered for sequences exhibiting ≥300 amino acid residues, ≥50% identity, and a query start position <3 amino acid residues in NCBI BLAST output format. A tabular summary of the homology observed for each species was compiled (Supplemental data 3).

### Structural Analysis of Full-Length SBHV4-S9 and MHC Class I Peptides

Structural analysis of the full-length sequences of SBHV4-S9 (YP_004940188.1), HLA-A (AZU89775.1), the *Piliocolobus tephrosceles* MHC class I-like peptide (African origin; closest homolog identified by Spaln3[32]), and the *Scleropages formosus* MHC class I-like peptide (XP_029104321.1) was performed using AlphaFold 2.3.2 [21]. Full-length sequences were employed to explore potential functional similarities beyond high-homology subregions. TM-scores for pairwise comparisons were normalized by the length of the shorter sequence. The resulting four PDB files were used to assess pairwise structural similarity via the TM-align web server [22], and visualized using ChimeraX software [23].

### Molecular Phylogenetic Analysis of MHC Class I-like and SBHV4-S9 Genes

To elucidate the molecular evolutionary history of SBHV4-S9 in relation to MHC class I-like genes, we performed DNA phylogenetic analysis.

The sequence corresponding to the high-homology region of SBHV4-S9 within HLA-A peptide was manually extracted from the Ensembl entry ENST00000706898.1 (HLA-A-216). For SBHV4-S9, the homologous region to HLA-A was extracted from the SBHV4 genome (NC_016448.1).

Subsequently, coding sequence data for all species available on the Ensembl FTP site (https://ftp.ensembl.org/pub/release-112) were downloaded and subjected to homology searches (-b 1 -d 1 -E 1e-10 -m BB). Resulting BLAST-formatted .txt files within this directory were filtered to retain sequences ≥900 bp in length, initiating with a query start position ≤15 (to ensure data integrity), yielding FASTA36 results for 126 animal species + 1 virus species (SBHV4-S9). All MHC class I-like sequences for these 126 animal species + 1 virus species (SBHV4-S9) were compiled into a single FASTA file.

To investigate associations with human variants, HLA-A haplotype sequences were collected from human genome CRAM files available from the International Genome Sample Resources (IGSR; https://www.internationalgenome.org/data-portal/data-collection/30x-grch38). A comprehensive list of available CRAM files (igsr_30x GRCh38.tsv.tsv) was downloaded from the IGSR website (Available data tab, as of July 2024). A list of up to four individuals per cohort with non-redundant Population designations was generated (igsr_30x GRCh38_rev.tsv; 104 individuals). CRAM files for these 104 individuals were retrieved. FASTQ files extracted from these CRAM files were mapped to HLA-A_gene_GRCh38.fa (EU445472.1). De novo assembly was performed on the resulting .sam files using SPAdes software (-k 31) [24]. Following careful examination of open reading frames and homologies, and excluding redundant HLA-A haplotypes, six human individuals exhibiting ≥99% homology to HLA-A CDS datum (GRCh38) were selected as candidates for phylogenetic analysis (IDs specified in Fig. 3). The FASTA data for these human individuals, previously recovered other species, 126 animal species including humans and SBHV4-S9 were concatenated into a final file (containing 133 fasta entries; Supplemental data 4). The presence of the motif characteristic of MHC class I molecules was confirmed across all FASTA data using MEME suite (https://meme-suite.org/meme/index.html) [25]. Finally, this dataset underwent global alignment with MAFFT [26] and trimming with ClipKIT [27], followed by ML phylogenetic tree inference using IQ-TREE 2 [28] (substitution model selection: -m MFP+LM; UFbootstrap and SHaLRT calculation). Branch support reliability was assessed using a composite S-value, calculated as the geometric mean of SHaLRT and UFbootstrap values output by IQ-TREE 2.

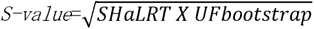

Nodes with S-values ≥87% were considered reliably supported and visualized using FigTree (https://github.com/rambaut/figtree). Initially, a cladogram was generated to illustrate branching patterns. Given the close phylogenetic affinity of SBHV4-S9 to primates among vertebrates, as of June 2025 we downloaded all primate genomes available in NCBI. Using the HLA-A peptide sequences of the SBHV4-S9 homologous region, we retrieved the corresponding coding sequences via Spaln3 (-H 1000). Species with excessive ambiguous nucleotides (N), extremely short nucleotide sequences, or in-frame stop codons were excluded, yielding a final dataset of 186 primate sequences with SBHV4-S9 (Supplemental data 5). The presence of an MHC class I motif in all sequences was verified. Following global alignment with MAFFT, phylogenetic inference was performed using IQ-TREE 2.

### Divergence Time Estimation of SBHV4-S9

From the final FASTA file used for ML analysis (Supplemental data 5, excluding SBHV4-S9), FASTA data for primates and outgroups (*Monodelphis domestica*, *Cavia porcellus*, *Ornithorhynchus anatinus*, *Oryctolagus cuniculus*) were constructed. Following global alignment with MAFFT and parameter acquisition with IQ-TREE 2, an XML file was generated using BEAUti [29] and BEAST 2 was executed [29] (ESS values ≥200 for all parameters). From the resulting treefile, a Maximum Clade Credibility tree was generated after discarding the initial 10% burn-in using TreeAnnotator [29]. Numerical values pertaining to the divergence time estimation between *P. tephrosceles* and 10 Old World monkeys were extracted from this Nexus format file.

### dN/dS Calculation

dN/dS (ω) values were calculated using CODEML in PAML [30]. For codon alignment, the FASTA file from Supplemental data 5 was aligned using PRANK software [31], employing the phylogenetic tree from Fig 4. To assess the selective pressure on the SBHV4-S9 gene, a branch-specific model (model = 2, NSsites = 0) was used to perform a likelihood ratio test (LRT) (Supplemental data 6). A null model was configured to fix the ω value for the SBHV4-S9 branch at 1 (fix_omega = 1, omega = 1.0), representing neutral evolution, while the ω values for all other branches were freely estimated. This was compared to an alternative model where the ω values for all branches, including SBHV4-S9, were allowed to be freely estimated (fix_omega = 0). The LRT was used to determine whether the ω value for the SBHV4-S9 branch was significantly less than 1.

### Molecular Phylogenetic Analysis of SLAMF6-like and Primate SBHV4-S1 Genes

To elucidate the co-evolutionary history of SBHV4-S1 with primate SLAMF6-like genes, the corresponding DNA sequences for the high-homology region between SBHV4-S1 and human SLAMF6 (AB590474.1) were obtained using human SLAMF6 peptides (Spaln3; -H 800). Species with excessive ambiguous nucleotides (N), extremely short nucleotide sequences, or in-frame stop codons were excluded, yielding SLAMF6-like sequences from 112 primate sequences. To assess preliminary orthology, motif searches were performed using MEME suite [25], as previously described. All sequences commonly exhibited Ig_3 motif characteristic of SLAMF6. Following global alignment with MAFFT, ML phylogenetic inference was conducted using IQ-TREE 2, employing -m MFP for substitution model selection, with Ultra-fast bootstrap and SHaLRT. The results were converted to the aforementioned S-values and visualized.

### Phylogenetic Relationship Between SBHV4 and Primates

We retrieved the complete set of 141 peptide sequences encoded by SBHV4 (NC_016448). Using these sequences, we queried the genomes of 244 primate species with Spaln3 (-H 100). Regions with at least one match were detected in 243 species. Among the SBHV4 genes, eight (S1, S8, S9, S11, S28, UL54, UL116, and A23) exhibited homology to primate genomes (Supplemental data 7). None of the primate species contained all eight genes. Homologous sequences for these eight SBHV4 genes were extracted in multi-FASTA format using the GFF files generated by Spaln3 for each primate species. After excluding one species lacking any homologous genes (*Tarsius wallacei*) and two species whose retrieved sequences contained excessive ambiguous nucleotides (N), the dataset comprised 241 primate species plus the eight SBHV4 genes (Supplemental data 8). These were used for the subsequent analyses.

To analyze the phylogenetic relationships between SBHV4 genes and 241 primate species (not all of which possess homologs for every SBHV4 gene), a hybrid alignment-free framework integrating syncmers [33], principal component analysis (PCA), and the Wasserstein distance (Earth Mover’s Distance; EMD) pipeline was developed (Fig. 7) (Code 2; The script implements the alignment-free phylogenetic analysis pipeline, which incorporates PCA-based selection of representative syncmers, fuzzy mapping using weighted Hamming distance, and the calculation of Optimal Transport (Wasserstein distance). Based on specified parameters, the script outputs the final frequency vectors, the original distance matrix, and Bootstrap distance matrices derived from k-mer level resampling.). The adopted Wasserstein distance is a statistical metric that quantifies the dissimilarity between two probability distributions and can be interpreted as the minimal amount of work required to transform the mass of one distribution into the shape of another. In previous studies, the Optimal Transport (OT) distance, specifically the Wasserstein distance, has been employed in single-cell analysis to measure the affinity or similarity between cellular gene expression profiles [34]. In this study, we adapted this approach by collecting multiple SBHV4 genes and their corresponding primate homologs. We utilized the frequency vectors, extracted via syncmers, and their subsequent PCA loadings to construct a high-dimensional, statistical frequency distribution representing each species, and to choose representative syncmers for the distance measurement among SBHV4 genes and primate homologues. A variant of the Wasserstein distance measurement (the Python *ot* library) was applied to quantify the similarity among the SBHV4 genes and their homologs, thereby aiming to elucidate the overall phylogenetic relationship between SBHV4 and primate species.

Owing to the OT property, the method enables overall differences in distributional shape, derived from the integrated information of multiple SBHV4 genes and their homologs in 241 primates, to be captured quantitatively, going beyond simple comparisons of means or modes. In contrast to traditional phylogenetic approaches based on nucleotide-sequence alignments, where distances are calculated from direct sequence similarity, the present method represents each species as a single, multi-variate statistical distribution by integrating the PCA-transformed syncmer feature space across multiple genes. Consequently, subtle shifts and deformations in distributional structure are reflected in the resulting distances, potentially capturing phylogenetic relationships grounded in a more holistic view of biological variation.

This integrative representation further confers robustness to missing data. The statistical distribution is defined by the aggregated features of all available SBHV4 homologs. Even when certain homologs are absent in a given species, the remaining homologs still contribute to the formation of that species’ multi-variate distribution, effectively diluting the impact of the missing information on the overall shape of the species’ feature space. This allows distances to be computed without forfeiting phylogenetic signal.

Unique canonical syncmers (70 bp) were first selected from the genes and homologs in each primate species and SBHV4. The overall process integrates feature selection (PCA) and final distribution construction.

First, for feature selection: To determine the most representative syncmers, initial frequency vectors were constructed by mapping all k-mers from each species onto the canonical syncmers. PCA was then performed on these initial frequency vectors, and 10,000 representative PCA-syncmers were chosen by feature selection based on PCA loadings. This procedure constituted a dimensionality-reduction step intended to mitigate computational burden. This process generates the input for the PCA-syncmer frequency vectors (matrix), which is detailed in the ‘Third’ section below.

Second, for the Cost Matrix: For the selected PCA-syncmers, a weighted Hamming distance was calculated by assigning a weight of 0.5 to transitions and 1.0 to transversions. The resulting 10,000 × 10,000 ground distance matrix was used as the cost matrix input for Wasserstein distance computation.

Third, for the Final Frequency Vectors (Mass Distribution): To construct the final statistical distribution, the k-mer sets derived from each species were treated as the query mass, which was then mapped onto the feature space defined by the representative PCA-syncmers (the reference feature space). The mapping process utilizes the weighted Hamming distances as follows; Weighted Hamming distances were measured between all k-mers from each primate species (query) and the 10,000 PCA-syncmers (reference), and nucleotide similarity was computed as “1.0 − (weighted Hamming distance)/(total nucleotides)”. Fuzzy mapping was performed using a similarity threshold of 0.85 (85%). When a mapping succeeded, the corresponding PCA-syncmer count was incremented with a weight equal to the observed similarity (e.g., 0.98). After aggregation and L1-normalization, the final PCA-syncmer frequency vectors were obtained for all primate species and SBHV4 and subsequently subjected to Wasserstein distance computation. Wasserstein distances were computed using the ot library implemented in Python.

Importantly, these parameters were not arbitrarily selected; rather, they were evaluated and adjusted in the context of the benchmarking analysis employing 15 mitochondrial genes from 49 primate species. In this analysis, phylogenetic trees reconstructed from Wasserstein distance matrices were compared to those derived using ML methods, which are conventionally employed for nucleotide alignment-based phylogenetics. Threshold values for fuzzy mapping and corrected counts for final frequency vectors to Wasserstein distance computation in 0.95 identity yielded topologies most similar to the 49 primate species phylogenetic tree using 15 mitochondrial genes [35, 36] and were therefore adopted for the primary analysis. In the SBHV4 analysis, however, the threshold of 0.85 was applied because the comparison involved viral genes and their primate homologs, as 0.95 identity failed to map the SBHV4 genes to the representative PCA-syncmers due to elevated sequence divergence. The mapping rates of SBHV4 to PCA-syncmers were 0%, 0.31%, 2.34%, and 11.46% at base-pair identity thresholds of 95%, 93%, 90%, and 85%, respectively. Since a clear, universal criterion for the mapping rate to be passed to the Wasserstein distance matrix does not exist, this study selected the optimal threshold based on a balance between Wasserstein distance stability (Coefficient of Variation, CV) and data coverage (information content). First, we performed a bootstrap trial with k-mer level resampling in each primates and evaluated this stability using the CV at base-pair identity thresholds of 93%, 90%, and 85%. The median CV values were 0.0145, 0.0141, and 0.0140, respectively. The proximity of these values indicated that computational stability was satisfied across all tested thresholds. A Bonferroni-corrected Dunn’s Post-hoc test detected a statistically significant difference (P < 0.05) between the threshold with the highest median CV (93% base-pair identity) and the 90% and 85% thresholds. Despite this statistical difference, which represents a minimal practical variation (a maximum difference of 0.0005 in the median CV), we proceeded to optimize based on data coverage. Thresholds with a mapping rate of less than 3% (93% and 90% base-pair identity) result in extreme sparsity (too few non-zero elements) in a distance vector. Even accounting for the genetic divergence between viruses and primates, such sparsity increases the risk of vulnerability due to information imbalance in phylogenetic analysis. Therefore, based on the criteria of maintaining the Wasserstein distance stability while minimizing the loss of feature information and maximizing phylogenetic discriminative power, the 85% base-pair identity threshold (mapping rate 11.46%) was selected as the optimal threshold. At this parameter (85% identity), the average mapping rate among all 241 primates and SBHV4 was 51.38% (93%: 39.12%, 90%: 42.43%).

Benchmark tests based on mitochondrial genes were conducted without inclusion of SBHV4 genes. For the benchmarking analysis, 50,000 PCA-syncmers were employed, the nucleotide-similarity threshold was set to 0.95, and all other parameters were maintained as described above. For the phylogenetic analysis encompassing SBHV4 and 241 primate species, the number of PCA-syncmers was reduced from the benchmark of 50,000 to 10,000. This reduction was necessitated by computational resource limitations stemming from the increased computational load of the Wasserstein distance matrix calculation, which was exacerbated by the inherent complexity between the viral sequence and its primate homologs.

### Benchmarking Against Maximum-Likelihood Test

Benchmarking was conducted using 49 primate species selected from the SBHV4 homology list:

- Apes: Family Hominidae and Family Hylobatidae (19 species)
- Tarsiers: Family Tarsiidae (3 species + Tarsius wallacei)
- New World monkeys: Family Aotidae (5 species) and Genus Saimiri (6 species)
- Old World monkeys: Genus Macaca (15 species)

From the mitochondrial genome of *Carlito syrichta* (NC_012774), all 13 protein-coding genes and two rRNAs were retrieved. Using fasta36 (-E 1e-10), 15 homologous DNA sequences were obtained for each of the 49 species. Multiple sequence alignments were performed gene-by-gene with mafft, then concatenated per species. IQ-TREE 2 was run with 15 partitions (-m MFP -alrt 10000 -bb 10000 -ninit 10000 -ntop 1000 -o Carlito_syrichta), producing a Q-matrix that was then applied in the Wasserstein distance analysis described above.

## Data availability

All data utilized in this study are presented within the Figures or as Supplemental Data.

## Acknowledgements

This work was partially supported by Grants-in-Aid for Scientific Research from the JSPS (24K21913 to E.H.). Computations were performed on the NIG supercomputer at ROIS National Institute of Genetics, Japan.

## Notes

### Competing Interest Statement

The authors have declared no competing interest.

### Summary of Updates

Major modification of the main manuscript text, Figure 7 and 9, and Python codes. 2 codes removed and 1 code (Code_2.txt) was largely modified, that is, the parameter settings were made more flexible.

